# DeepAIR: a deep-learning framework for effective integration of sequence and 3D structure to enable adaptive immune receptor analysis

**DOI:** 10.1101/2022.09.30.510251

**Authors:** Yu Zhao, Bing He, Chen Li, Zhimeng Xu, Xiaona Su, Jamie Rossjohn, Jiangning Song, Jianhua Yao

## Abstract

Structural docking between the adaptive immune receptors (AIRs), including T cell receptors (TCRs) and B cell receptors (BCRs), and their cognate antigens is one of the most fundamental processes in adaptive immunity. However, current methods for predicting AIR-antigen binding largely rely on sequence-derived features of AIRs, omitting the structure features that are essential for binding affinity. In this study, we present a deep-learning framework, termed DeepAIR, for the accurate prediction of AIR-antigen binding by integrating both sequence and structure features of AIRs. DeepAIR consists of three feature encoders (a trainable-embedding-layer-based gene encoder, a transformer-based sequence encoder, and a pre-trained AlphaFold2-based structure encoder), a gating-based attention mechanism to extract important features, and a tensor fusion mechanism to integrate obtained features. We train and evaluate DeepAIR on three downstream prediction tasks, including the prediction of AIR-antigen binding affinity, the prediction of AIR-antigen binding reactivity, and the classification of the immune repertoire. On five representative datasets, DeepAIR shows outstanding prediction performance in terms of AUC (area under the ROC curve) in predicting the binding reactivity to various antigens, as well as the classification of immune repertoire for nasopharyngeal carcinoma (NPC) and inflammatory bowel disease (IBD). DeepAIR is freely available for academic purposes at https://github.com/TencentAILabHealthcare/DeepAIR. We anticipate that DeepAIR can serve as a useful tool for characterizing and profiling antigen binding AIRs, thereby informing the design of personalized immunotherapy.

**Highlights:**

1. Integrating predicted AIR structures using AlphaFold2 significantly improves the prediction accuracy of the binding reactivity between AIRs and antigens.
2. DeepAIR is featured by a novel deep learning architecture that leverages both the gating-based attention mechanism and tensor fusion mechanism to effectively extract and integrate informative features from three feature encoders, including a trainable embedding-layer-based gene encoder, a transformer-based sequence encoder, and a pre-trained AlphaFold2-based structure encoder.
3. DeepAIR is implemented as a biologically interpretable deep learning framework that highlights the key residues in both α and β chains that are critical for predicting the AIR-antigen binding.

## Introduction

Adaptive immune receptors (AIRs) recognize antigens to activate the ensuing immune responses, thereby scrutinizing and killing the tumor cells and invading pathogens^1^. T cell receptor (TCR) and B cell receptor (BCR) are two major types of AIRs. TCRs bind to the peptides (antigens) presented by the major histocompatibility complex (i.e., peptide-MHC, pMHC) on the cell surface^2^, while BCRs directly recognize native and cognate antigens^3^. Both TCR and BCR are composed of two polypeptide chains (i.e., alpha-beta or light-heavy) that form three-dimensional structures of the complementarity-determining region (CDR) loops (i.e., CDR1, CDR2, and CDR3) to bind the antigen^4^. The CDR1 and CDR2 loops of TCR often – but not always – contribute to MHC binding ^5^, while the CDR3 loops can play a prominent role in contacting the peptide, although CDR1/2 loops are known to mediate peptide contacts too^6^. For both TCR and BCR, the CDR3 loop is the most diverse region that has been widely used in the studies of immune repertoire^7–9^, which is defined as the sum of TCRs and BCRs that makes the organism’s adaptive immune system. Each chain is encoded by a somatically recombined gene sequence of the Variable (V) gene segments, the Diversity (D) gene segments (presented in half of the chains), and the Joining (J) gene segments. The genetic rearrangement of V(D)J gene segments generates a highly polymorphic AIR repertoire, which contains approximately 10^15^ to 10^61^ different receptors in human, allowing for the scrutinization and recognition of various antigens^10,11^. Accurate identification of the AIR-antigen recognition is therefore crucially important for understanding the adaptive immune system and designing immunotherapies and vaccines.

High-throughput sequencing bulk techniques have been widely applied to profile the V(D)J genes and the clonal diversity of AIRs^12^. The availability of such sequence data of V(D)J genes has allowed for clustering of the AIRs that recognize the same antigen based on the sequence-derived features^13,14^. However, high-throughput bulk sequencing techniques often capture only one chain of AIR, which is insufficient to profile the complete sequence features of the receptor, thereby hindering the development of a reliable prediction model for the AIR-antigen recognition based on the sequence features^12^. Recent advances in single-cell immune repertoire sequencing technologies have enabled the capture of both chains of the receptor, providing complete V(D)J gene sequencing data for the construction of AIR-antigen binding prediction models, such as GLIPH^13^, TCRdist^15^, DeepTCR^7^, TCRAI^8^, and soNNia^9^.

Among these models, GLIPH and TCRdist are two traditional statistical approaches, while others leverage state-of-the-art deep-learning technologies. GLIPH clusters TCRs that are predicted to bind the same pMHC according to the local sequence motifs and the global similarity of CDR3 sequences, which is the most diverse region of the CDR^13^. In contrast, TCRdist uses a similarity-weighted Hamming distance to measure the sequence similarities between TCRs and accordingly is able to identify antigen binding TCRs^15^. DeepTCR predicts antigen-TCR binding reactivity and affinity by incorporating the features from CDR3 sequences and V(D)J gene usage with a variational autoencoder (VAE)^7^. Similar to DeepTCR, TCRAI deploys a deep convolutional network built from CDR3 sequences and V(D)J genes to predict the TCR binding reactivity^8^. soNNia integrates interpretable knowledge-based models of immune receptor generation with deep-learning approaches to characterize the TCR and BCR binding reactivity, respectively^9^. As expected, deep-learning-based models, such as DeepTCR^7^ and TCRAI^8^, usually demonstrated superior prediction performance than traditional statistical models, such as GLIPH^13^ and TCRdist^15^. It is also worth noticing that most of the methods were designed for the AIR-antigen binding reactivity of TCR only, whereas soNNia is the only currently available method for both TCR and BCR. All these methods only used sequence-derived features to construct the machine-learning models. However, the structures of AIR play fundamental roles in recognizing and interacting with the antigen^16,17^. Despite the shortage of structural data of AIR receptors due to the high experimental cost, a wealth of accurately predicted structural data of AIRs have been made available due to the recent breakthrough of protein structure predictor, AlphaFold2^18^. It is now possible to construct more advanced and accurate computational models for AIR-antigen binding prediction.

In this study, we present a deep learning framework, termed DeepAIR, for AIR-antigen binding analysis, The functionality of DeepAIR includes AIR-antigen binding prediction,, and immune repertoire classification. Using a specifically designed gating-based attention mechanism and a tensor fusion mechanism, DeepAIR leverages the AlphaFold2-predicted AIR structure information to make the AIR-antigen binding prediction. Our benchmarking experiments demonstrate that on five datasets harboring 136,253 AIRs and 13 antigens (Supplementary Table 1), DeepAIR achieved superior prediction performance in terms of AUC (area under the ROC curve) across all the three tasks of AIR-antigen analysis compared to state-of-the-art approaches, including TCRAI, DeepTCR, and soNNia (Table 1).

**Table 1.** Performance of different methods for AIR-antigen binding analysis using single-cell immune repertoire data.

| Methods | AIR |  | Structure Feature | Binding Affinity |  | Binding Reactivity |  | Immune Repertoire Classification |  |
| --- | --- | --- | --- | --- | --- | --- | --- | --- | --- |
|  |  |  |  |  |  |  |  | AUC<sup>3</sup> |  |
|  |  |  |  |  |  |  |  | MIL-Voting |  |
|  | TCR | BCR |  | Pearson's correlation | AUC | TCR AUC<sup>1</sup> | BCR AUC<sup>2</sup> | TCR | BCR |
| **DeepAIR** | √ | √ | √ | 0.813 | 0.912 | 0.942 | 0.913 | 1 | 1 |
| **DeepAIR-stru** | √ | √ | √ | 0.800 | 0.769 | 0.882 | 0.722 | / | / |
| **DeepAIR-seq** | √ | √ | / | 0.733 | 0.745 | 0.854 | 0.864 | / | / |
| **TCRAI** | √ | / | / | / | / | 0.761 | / | / | / |
| **DeepTCR** | √ | / | / | 0.754 | 0.876 | 0.820 | / | / | / |
| **soNNia** | √ | √ | / | / | / | 0.796 | 0.660 | / | / |

<sup>1</sup> median value of the area under the ROC curve (AUC) for predicting the TCR binding reactivity for seven pMHC multimers using the leave-one-out cross-validation strategy.
<sup>2</sup> median value of the AUC for predicting the BCR binding reactivity for 4 antigens.
<sup>3</sup> median value of the AUC for the classifications for NPC and IBD.

## Results

### DeepAIR is a deep-learning framework by integrating 3D structure information for AIR-antigen binding prediction

The CDR3 loop of AIR is the most diverse CDR loop that plays a prominent role in contacting the epitope in the AIR-antigen binding complex^6,14^. Thus, the information of CDR3 sequence was widely used in previous methods, such as DeepTCR^8^ and TCRAI^17^, for the prediction of TCR-pMHC binding. We hypothesize that the structure of the CDR3 region is important for constructing an accurate model for AIR-antigen binding prediction. To examine this, we collected experimentally validated structures of two TCR-pMHC binding complexes (PDB ID: 1OGA^19^ and 3HG1^20^) from the PDB database^21^ (Figure 1A). It is known that the binding site (contact residue) mediates the binding between the AIR and the antigen^22^. Figure 1A illustrates the binding sites located on different chains of two TCRs according to the structures. We also collected the TCR sequences that bind to the same epitopes from the 10x website^23^ (Supplementary Table 1). From the collected sequences, we found it challenging to identify the binding sites from these sequences as the motifs containing the binding sites are not distinguishable (Figure 1A), which explains the difficulty of predicting the binding affinity using AIR sequences^22^. Beyond that, structure data is more distinguishable than sequence data in determining the binding reactivity of the AIR-antigen (Figure 1B-1C), e.g., for TCRs bind to the epitope YLQPRTFLL (human leukocyte antigen allele: HLA-A0201) of SARS-CoV-2 virus, although their CDR3 sequences are substituted 1∼5 amino acids, their CDR3 structures are nearly the same. (Figure 1B). But for AIRs bind to different epitopes LTDEMIAQY(HLA-A0101) and TTDPSFLGRY(HLA-A0201), respectively, of SARS-CoV-2 virus^23^, we found that their CDR3 structures show a larger difference than the sequences (Figure 1C). The above observations suggest that incorporating the structure information of the CDR3 region into the DeepAIR model might help boost the prediction performance.

**Figure 1.**
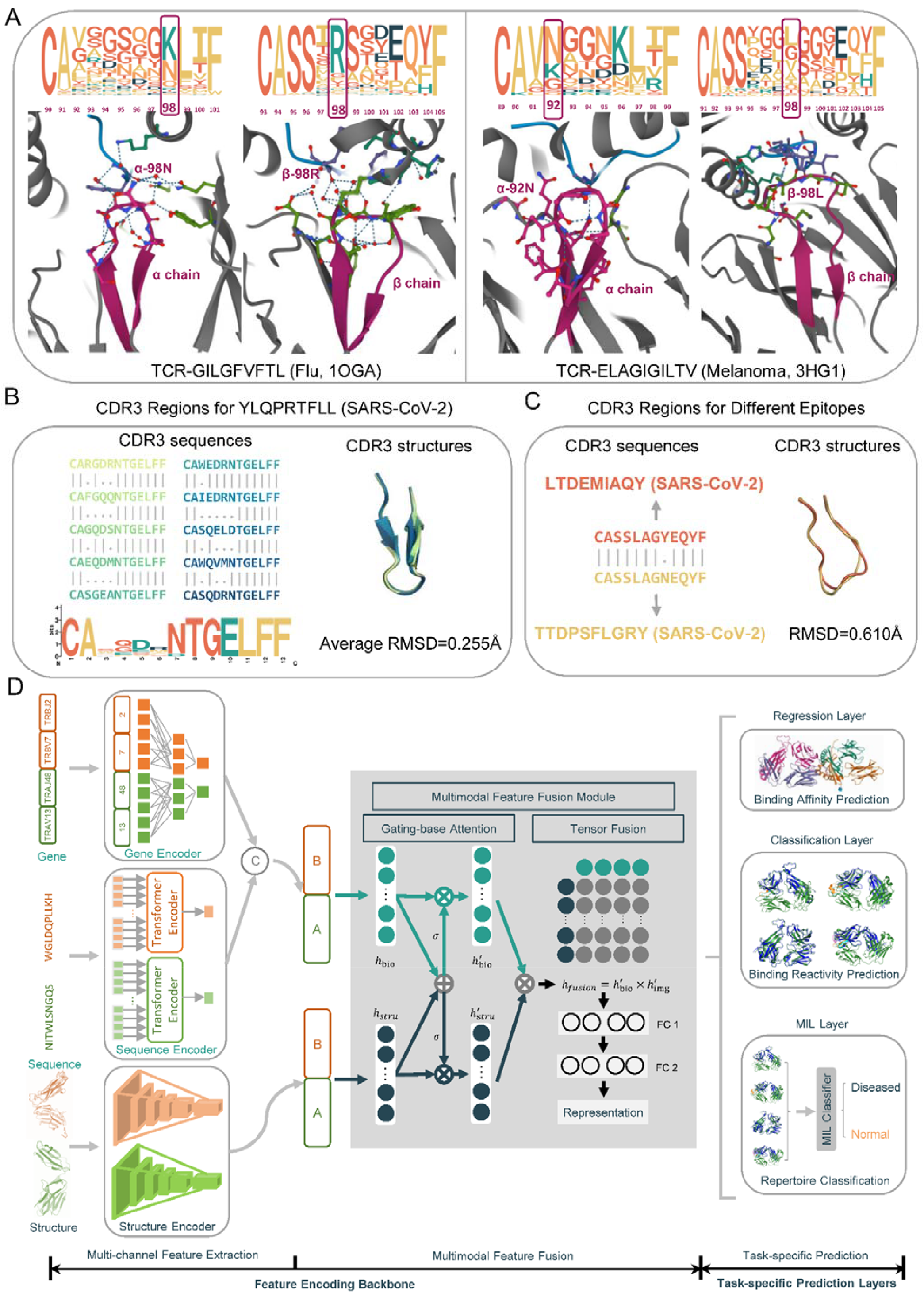
Constructing the computational framework of DeepAIR. (**A**) It remains a challenge to identify the binding site (contact residue) from sequence information. According to the sequence logo (top), the sequences are not distinguishable for the binding site discovered in the crystal structure (bottom) of the CDR3 region of the TCR α and β chains in the TCR-GILGFVFTL (Flu, PDB ID: 1OGA) binding complex (left) and TCR-ELAGIGILTV (Melanoma, PDB ID: 3HG1) binding complex (right), respectively. The sequence logos were rendered using TCRs from the 10x dataset (Supplementary Table 1). The size of the amino acid in the sequence logo indicates the frequency of the amino acid at the corresponding position. The position of the binding site (contact residue) is highlighted with a frame. (**B**) Sequences (left) of the CDR3 region that bind to the “YLQPRTFLL” from the SARS-CoV-2 virus are less stable than the structures (right). These 10 CDR3 sequences vary considerably but are structurally similar (RMSD=0.255 Å). The sequence and structure of the same TCR are labeled by the same color. (**C**) It is relatively easy and straightforward to distinguish the CDR3 regions that recognizing different antigens using the structures (right) than sequences (left). The sequence and structure of the CDR3 for binding to “LTDEMIAQY” (SARS-CoV-2) are colored in orange, while that for “TTDPSFLGRY” (SARS-CoV-2) are colored in yellow. The two sequences have only one amino acid difference; however, the CDR3 structures have a larger difference (RMSD=0.610 Å). The sequences in the panels (B) and (C) are retrieved from the SARS-CoV-2 virus dataset (Supplementary Table 1). (**D**) Flowchart of DeepAIR. There are three major processing stages in DeepAIR, including multi-channel feature extraction, multimodal feature fusion, and task-specific prediction. At the multi-channel feature extraction stage, three feature encoders are involved and used to extract informative features from the gene, sequence, and structure inputs. Then the resulting features produced by three different encoders are further integrated via a gating-based attention mechanism as well as the tensor fusion at the multimodal feature fusion stage to generate a comprehensive representation. Finally, at the task-specific prediction stage, specifically designed prediction layers are utilized to map the obtained representations to the output results.

DeepAIR has three major processing stages, including multi-channel feature extraction, multimodal feature fusion, and task-specific prediction (Figure 1D). In the multi-channel feature extraction stage, DeepAIR utilizes three feature encoders including the gene encoder, sequence encoder, and structure encoder to comprehensively encode the AIR. Specifically, the gene encoder embeds the V(D)J gene usage information via a trainable embedding layer. The sequence encoder generates a high-level representation of sequence information for the paired chains based on a multi-layer Transformer^20^. The structure encoder utilizes the pre-trained AlphaFold2 to extract the initial structure-information embedded features and then employs concatenated 1D convolutional layers to fine-tune and recalibrate these features specifically for the down-steam adaptive immune receptor analysis tasks. Then, in the multimodal feature fusion stage, a fusion module extracts key features from the obtained structure, sequence, and gene embeddings via a gating-based attention mechanism and then integrates them with the tensor fusion mechanism. Finally, in the task-specific prediction stage, the integrated features are fed into the task-specific prediction layers for the downstream AIR-antigen analysis tasks, including predicting the binding affinity with the regression layer, predicting the binding reactivity with the classification layer, and conducting the immune repertoire classification via the multi-instance learning (MIL) layer (Figure 1D).

To objectively characterize the contribution of the structure information, we created two variants of DeepAIR, namely DeepAIR-stru, and DeepAIR-seq. DeepAIR-stru is a model that uses only the structure information while DeepAIR-seq is a model that learns from sequence and the V(D)J gene usage information.

### The way of DeepAIR to obtain the AIR structure

Compared to over 277 million TCRs and BCRs with known sequences in the TCRdb^24^ database and Immune Epitope Database (IEDB) database^24^, there are only 858 experimentally validated structures available for human TCRs and 3,333 experimentally validated structures available for human BCRs in the PDB database^21^. Due to the limited availability of most AIR structures, we employed AlphaFold2 to predict the unliganded AIR structure and construct the DeepAIR model. The accuracy of the predicted AIR structure is therefore important for the prediction performance of DeepAIR. To find the best way of predicting the AIR structure using AlphaFold2, we collected experimentally validated structures of TCRs and BCRs with and without the antigen binding from the PDB database^21^. Then we predicted the AIR structure with AlphaFold2 using the amino acid sequences of the full-length beta/heavy chains and that of the CDR3 regions (Figure 2A-2D), respectively. The prediction accuracy was measured using the root mean square deviation (RMSD) between the predicted and the experimentally validated AIR structures.

**Figure 2.**
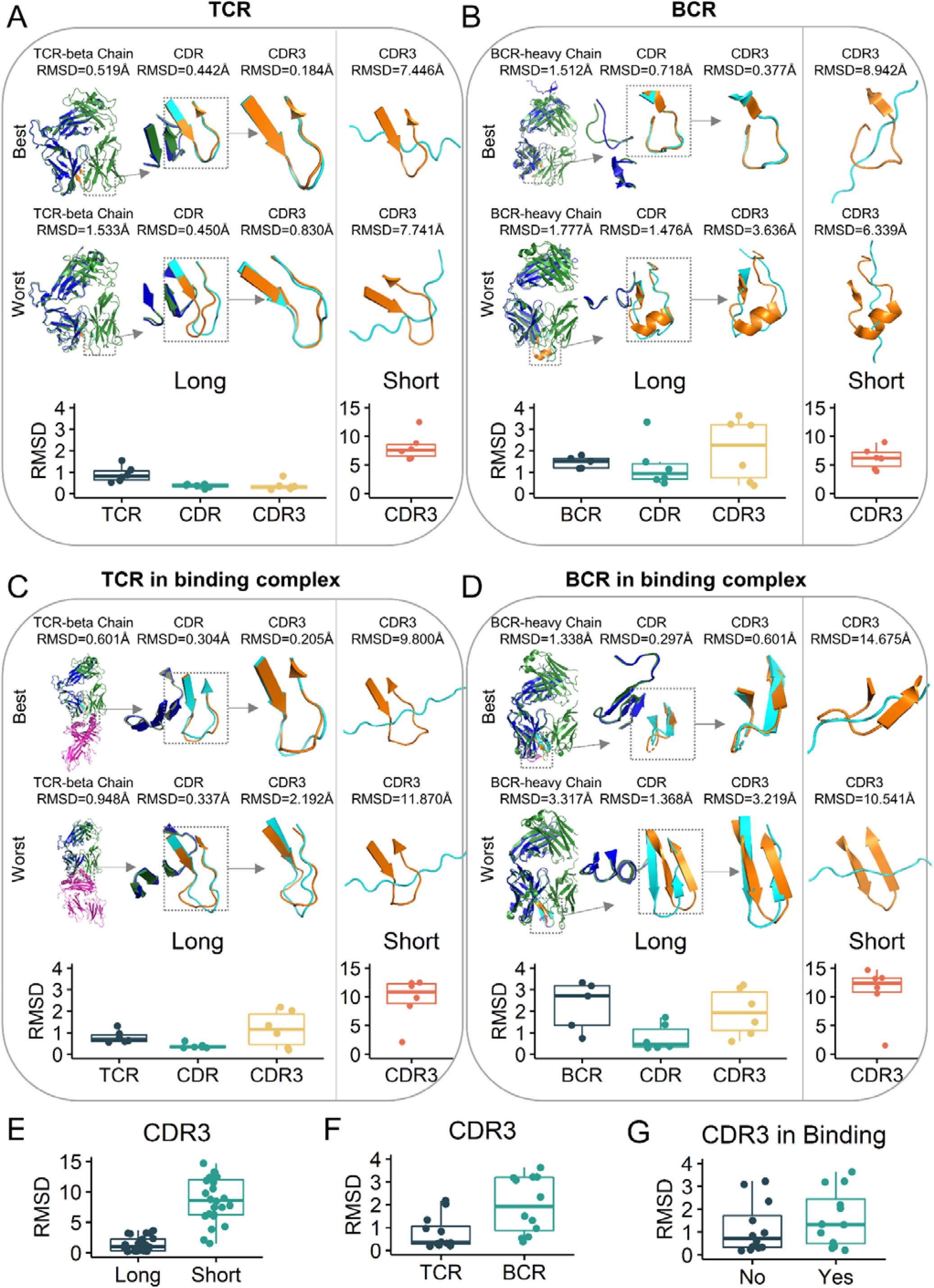
Evaluating the predicted AIR structures using AlphaFold2. The comparison between the predicted structure (blue) and the experimentally validated structure (orange) for (**A**) TCR, (**B**) BCR, (**C**) TCR in the binding complex, and (**D**) BCR in the binding complex. For each comparison, there were six predicted structures and six experimentally validated structures (Supplementary Table 2 and Supplementary Table 3). The root mean square deviation (RMSD) was used to measure the difference between the predicted and experimentally validated structures. The structures were predicted using the full AIR beta/heavy chain sequence (long sequence) and CDR3 sequence (short sequence), respectively. The structures from the prediction with the lowest (best) and highest (worst) RMSD of CDR3 are above the boxplot of the RMSD values. For each line, from left to right, there are structures of the full TCR beta chain (BCR heavy chain), CDR, and CDR3 from the prediction using the long sequence, and the structure of CDR3 from the prediction using the short sequence. For the prediction using the long sequence, RMSD was measured for the full TCR beta chain (BCR heavy chain), CDR, and CDR3, respectively. (**E**) The boxplot of RMSD for the predicted CDR3 using the long sequence and short sequence, respectively. (**F**) The boxplot of RMSD for the predicted CDR3 from TCR and BCR, respectively. (**G**) The boxplot of RMSD between the predicted CDR3 and the experimentally validated CDR3 regions with and without antigen binding, respectively.

In particular, we focused on the prediction accuracy of the CDR3 loop, which is the most diverse part of the AIR structure. The predicted CDR3 structures using the sequences of the full-length beta/heavy chains (long sequence, median RMSD=0.964 Å) had much lower RMSD values than that using only the sequences of the CDR3 regions (short sequence, median RMSD=8.598 Å) (Figure 2E, Supplementary Table 2 and Supplementary Table 3). The median RMSD for the predicted CDR3 structures using the long sequences was similar to that reported by AlphaFold2 on the CASP14 dataset^21^. Therefore, we used the long sequences to predict the AIR structures for quantifying the predictive capability of DeepAIR.

Interestingly, the predicted TCR structures appeared to be more accurate than the predicted BCR structures (Figure 2F). The median RMSD values for the predicted CDR3 structures of TCR and BCR using long sequences were 0.35 Å and 1.92 Å, respectively (Supplementary Table 2 and Supplementary Table 3). The results suggest that AlphaFold2 is not good at predicting the CDR3 structure of BCR. Moreover, the antigen binding decreased the prediction accuracy for the CDR3 structure (Figure 2G). The median RMSD values for the predicted CDR3 structures of AIRs compared to the experimentally validated CDR3 structures of AIRs with and without antigen binding were 1.42 Å and 0.46 Å, respectively (Supplementary Table 2 and Supplementary Table 3), suggesting that antigen binding may change the structure of CDR3 which can increase the difficulty of predicting the structure.

### Prediction of the AIR-antigen binding affinity

The antigen binding is based on the affinity between AIR and antigen. Currently, there is no reliable computational approach for predicting the exact binding affinity, especially for the TCR-pMHC binding^22^. In this study, we used the counts of unique TCR molecules that were captured by the pMHC as the observed proxy of AIR-antigen binding affinity^23^. We used the unique molecular identifier (UMI) to represent each unique TCR molecule. UMI is a type of molecular barcodes that provides error correction and increased accuracy during sequencing. These molecular barcodes are short sequences used to uniquely tag each molecule in a sample library. Due to the lack of BCR AIR-antigen binding affinity data, we instead focused on the prediction of TCR AIR-antigen binding affinity in this study.

We obtained the pMHC-captured single-cell TCR data from the 10x website^23^, which includes the single-cell TCRs captured by 44 pMHC multimers and six negative controls from four donors. The data was curated using the Integrative COntext-specific Normalization (ICON) workflow to remove the low-quality TCRs and false-positive bindings^22^. After data curation, 38,558 paired TCR α/β chains that bind to seven pMHC multimers, including ELAGIGILTV (HLA-A0201) from MART-1 protein of melanoma, GILGFVFTL (HLA-A0201) from M1 protein of influenza virus (flu), KLGGALQAK (HLA-A0301) from IE1 protein of cytomegalovirus (CMV), GLCTLVAML (HLA-A0201) from BMLF1 protein of Epstein-Barr virus (EBV), AVFDRKSDAK (HLA-A1101) from EBNA4 protein of EBV, IVTDFSVIK (HLA-A1101) from EBNA3B protein of EBV, and RAKFKQLL (HLA-B0801) from BZLF1 protein of EBV, were used in this study (Supplementary Table1).

The prediction of the AIR-antigen binding affinity is solved as a regression task in the DeepAIR framework. For each pMHC (antigen), its paired TCRs in the data set were randomly split into training data (70%), validation data (20%), and testing data (10%). The binding affinity prediction model was trained using the training data, optimized using the validation data, and tested independently using the independent test data. Since TCRAI and soNNia do not predict the biding affinity, we compared the performance of DeepAIR with that of DeepAIR-stru, DeepAIR-seq and DeepTCR. The affinities predicted by DeepAIR achieved the highest Pearson’s correlation with the experimentally observed proxy of binding affinities^19^ (Figure 3A). Meanwhile, DeepAIR achieved the lowest mean squared error (MSE) and mean absolute error (MAE) values, suggesting that the AIR-antigen affinities predicted by DeepAIR were the closest to the experimental observations (Figure 3C). Next, we examined whether the predicted binding affinity was accurate enough to determine the specific binding between the TCR and the pMHC. We used the receiver-operating characteristic (ROC) curve to illustrate the power of the predicted affinity in distinguishing the experimentally observed TCR-pMHC binding. The area under the ROC curve (AUC) is the aggregated measure of the performance for this task. As a result, DeepAIR achieved an AUC of 0.912, which was significantly better than that from any of the other methods (Figure 3B). It is also of particular interest to note that DeepAIR-stru outperformed DeepAIR-seq across all the comparisons (Figures 3A-3C), suggesting the contribution of the structure data to improve the prediction performance.

**Figure 3.**
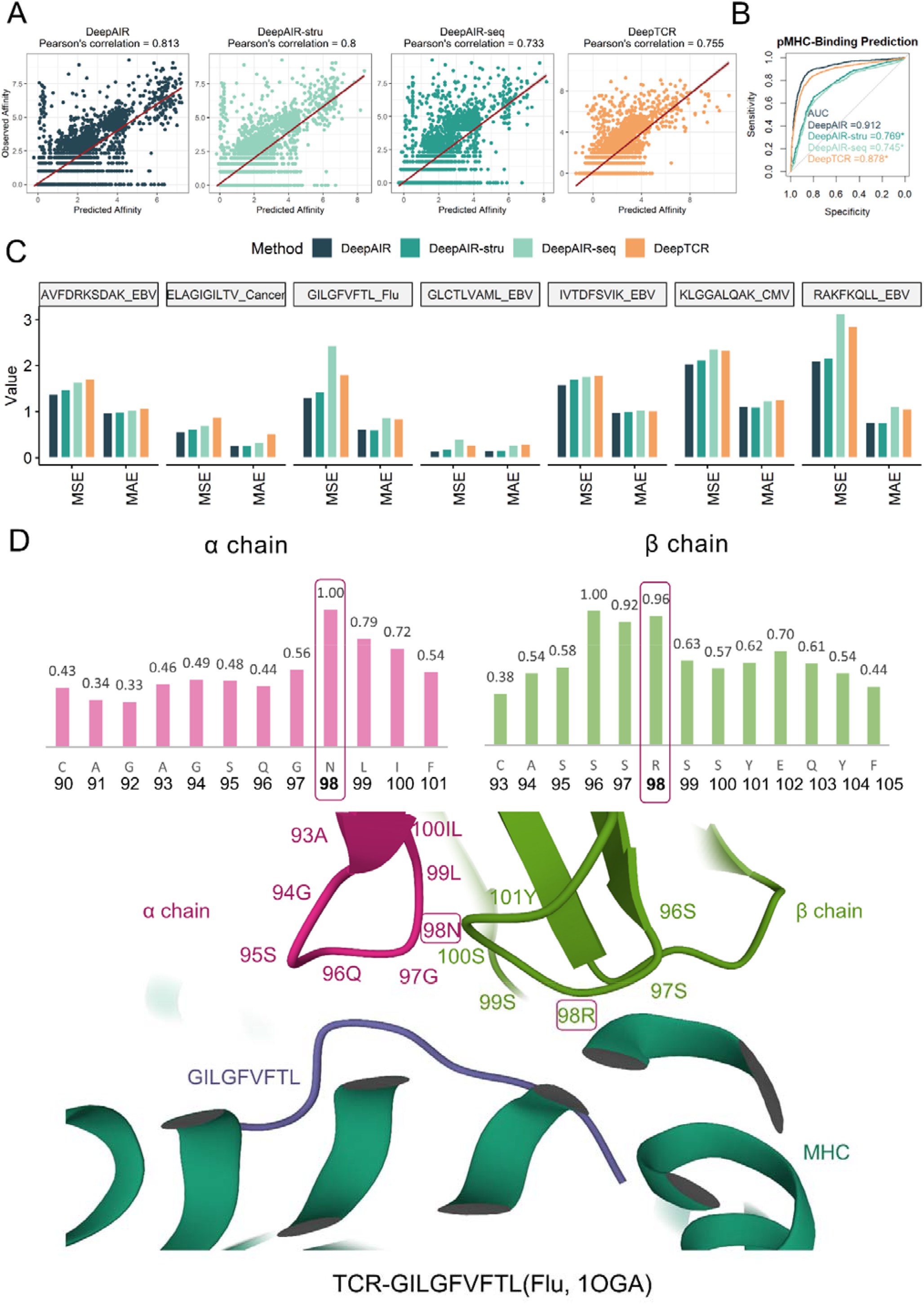
Performance comparison of DeepAIR and DeepTCR TCR binding affinity predictions. (**A**) Scatter plots demonstrating the correlations between experimentally validated AIR-antigen binding affinity values and the predicted binding affinity values by DeepAIR and DeepTCR respectively on the 7 pMHC multimer dataset. The line was generated to show the best fit using a linear regression model. (**B**) ROC curves for determining the experimentally observed pMHC-binding using predicted affinity. The *p*-value is produced by the comparison of ROC curves using the DeLong test. DeepAIR achieved statistically higher performance than the other three models. * means *p*-value<0.05 in the comparison with DeepAIR. (**C**) The mean squared error (MSE) and the mean absolute error (MAE) values between the predicted AIR-antigen binding affinity values and the experimentally validated affinity values for each pMHC multimer using DeepAIR and DeepTCR. (**D**) The normalized DeepAIR attention weights for each residue in the CDR3 region of α chain (up left) and β chain (up right), respectively, and the experimentally validated crystal structure (PDB ID: 1OGA) (bottom) of the TCR that binds to the GILGFVFTL (Flu, PDB ID: 1OGA). A higher attention weight indicates the residue is more important to the prediction of AIR-antigen binding affinity. The amino acids with high attention weight, such as α-98N and β-98R, are contact residues in the crystal structure. The α-98N stabilizes the TCR structure formed by the α chain and β chain, while β-98R stabilizes the binding between TCR and GILGFVFTL (Flu, PDB ID: 1OGA). The α-98N and β-98R residues are highlighted with a frame.

To better understand and interpret how well DeepAIR could predict the AIR-antigen binding affinity, we extracted the attention weights of every residue from the model that predicts the affinity to GILGFVFTL. A high weight indicates the residue is important to the prediction of AIR-antigen binding affinity. For example, according to the attention weight, the amino acid residue Arginine (Arg, R) at the β-98 position is crucial to the binding between TCR and HLA-A2-GILGFVFTL (M1 protein, flu) (Figure 3D). Then, we examined the crystal structure of the TCR-GILGFVFTL binding complex that was collected from PDB ID: 1OGA^19^. We note that β-98R is the contacting residue between the TCR-β chain and GILGFVFTL (Figure 3D). In this case, DeepAIR precisely captured the important part of the TCR that impacts the AIR-antigen binding. Moreover, we also note DeepAIR highlights the importance of Asparagine (Asn, N) at the α-98 position. This residue is the contact residue between the α chain and β chain that stabilizes the structure of TCR. The results indicate that DeepAIR learned that the stabilization of the paired α-β structure is important for the binding affinity between the TCR and antigen. Moreover, DeepAIR highlights similar partial structures in TCRs with high binding affinity but not in that with low binding affinity to GILGFVFTL (M1 protein, flu) (Supplementary Figure 1). Taken together, DeepAIR not only accurately predicts the AIR-antigen binding affinity, but also reveals the important residues that directly contribute to the binding of AIRs to the antigens.

### Prediction of the AIR-antigen binding reactivity

In addition to the use of the AIR-antigen binding affinity, a common strategy for predicting the AIR-antigen binding reactivity is to effectively learn the patterns from the AIRs that bind to the same antigen. This is considered and solved as a classification task in DeepAIR. To evaluate the performance of DeepAIR for predicting the binding reactivity of TCR, we collected experimentally validated pMHC-specific TCRs from various sources, including the 10x Genomics website^25^ and a SARS-CoV-2 virus study^26^. The 10x dataset, which has 38,558 paired TCR α/β chains that bind to 7 pMHC multimers, is the same one as we used for the AIR-antigen binding affinity prediction. The SARS-CoV-2 virus dataset has 592 paired TCR α/β chains that bind to 3 pMHC multimers from the SARS-CoV-2 virus. These pMHC multimers include LTDEMIAQY (HLA-A0101) and YLQPRTFLL (HLA-A0201) from the spike protein, and TTDPSFLGRY (HLA-A0201) from the ORF1ab polyprotein. Therefore, a total of 39,150 TCRs for 10 pMHC multimers were used in the prediction of binding reactivity.

To investigate whether the deep-learning model can predict the AIR-antigen binding reactivity for the unseen TCRs, we randomly split the TCRs into the training data (70%), validation data (20%), and testing data (10%) as we did in the binding affinity prediction task. DeepAIR achieved a median AUC of 0.904 in predicting the AIR-antigen binding reactivity for the 10 pMHC multimers (Supplementary Table 4), significantly outperforming all the other methods, including DeepAIR-stru (median AUC=0.873), DeepAIR-seq (median AUC=0.827), TCRAI (median AUC=0. 843), DeepTCR (median AUC=0.846), and soNNia (median AUC=0. 782) (Figure 4A and Supplementary Table 4). Interestingly, the majority of the methods achieved better performance in predicting the TCRs that specifically bind to ELAGIGILTV (MART-1 protein from Melanoma), and worse performance in predicting TCRs that specifically bind to LTDEMIAQY (spike protein from SARS-CoV-2 virus). These results suggest that TCRs for LTDEMIAQY (the spike protein from the SARS-CoV-2 virus) is more diverse than that for ELAGIGILTV (the MART-1 protein from Melanoma).

**Figure 4.**
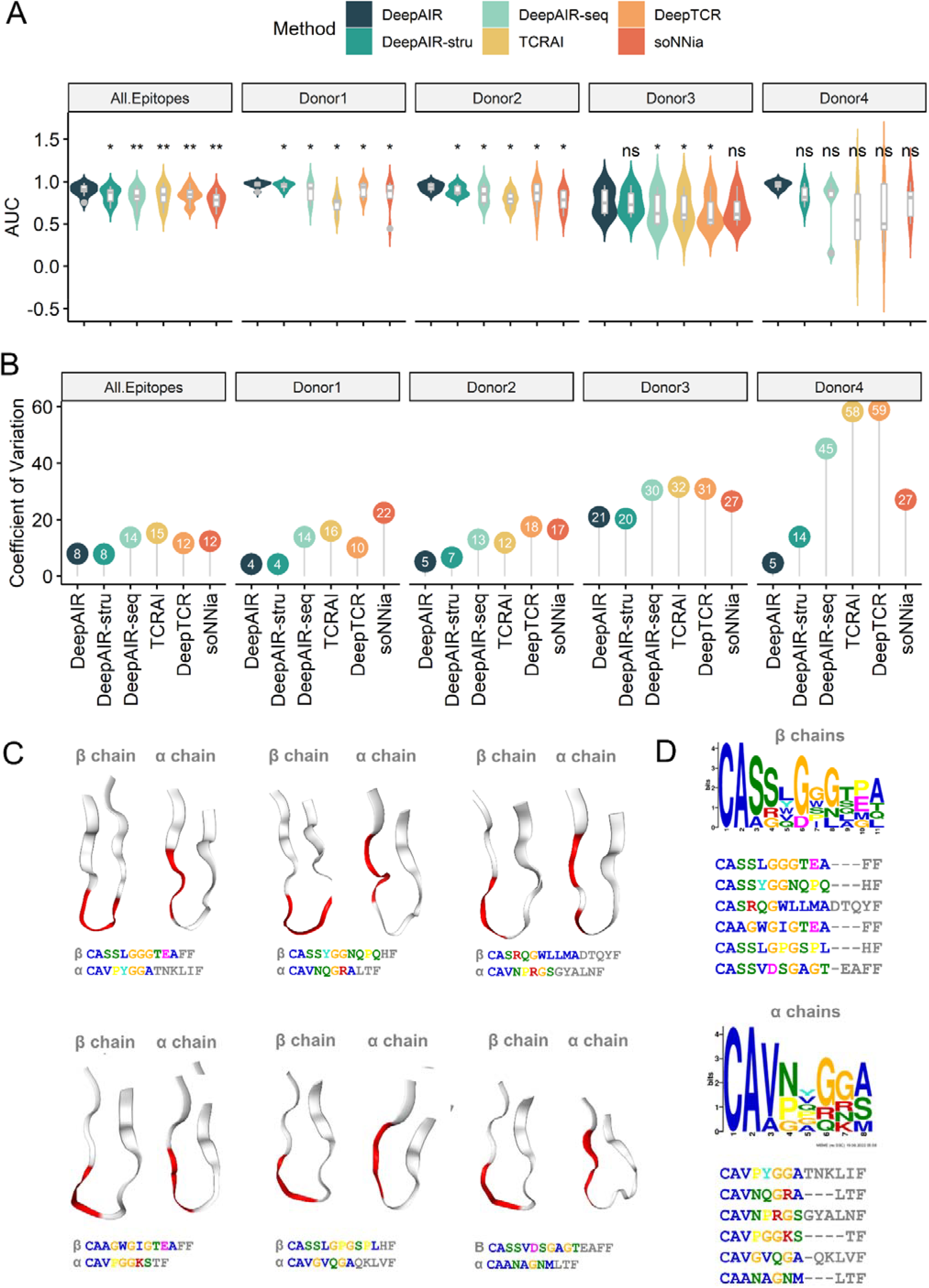
Performance comparison between DeepAIR and state-of-the-art approaches for AIR-antigen binding reactivity prediction. (**A**) Violin plots of the area under the ROC curve (AUC) values for DeepAIR and state-of-the-art approaches in predicting the binding reactivity. The “All Epitopes” subplot includes 10 pMHC multimers, 7 of which are from the 10x dataset and the rest 3 multimers are from the SARS-CoV-2 virus study (Supplementary Table 1). The AUC for “All Epitopes” measures the performance of the obtained model on the test data. The “Donor1”, “Donor2”, “Donor3” and “Donor4” show the AUC values obtained in the leave-one-out test, where the donor data was used for testing the performance. ‘ns’ means not significant, * means *p*-value<0.05, and ** means *p*-value<0.01 in the comparison with DeepAIR. (**B**) The coefficient of variance for AUC performance of DeepAIR and the compared methods for all epitopes and each donor, respectively. (**C**) Structure patterns learned by DeepAIR for predicting the antigen binding TCRs. The structures of TCRs with specific binding to ELAGIGILTV (MART-1 protein, melanoma) are illustrated, and the most salient regions supporting DeepAIR to make the prediction are highlighted in red. (**D**) The sequences and recognized motifs of the corresponding beta chains and alpha chains of TCRs with specific binding to ELAGIGILTV (MART-1 protein, melanoma) in subfigure (C). The sample ELAGIGILTV-specific TCR pairs are quite diverse in the sequence view.

In practical use, a common scenario is that we use a well-trained model to predict the AIR-antigen binding reactivity for the TCRs from individuals independent from the training cohort. To evaluate the model performance in this scenario, we performed the leave-one-out cross-validation. There are four donors in the 10x dataset. Donor 1 and donor 2 have 7,280 and 26,705 TCRs, respectively, which bind to 7 pMHC multimers. Donor 3 has 2,040 TCRs that bind to 6 pMHC multimers. Donor 4 has 2,533 TCRs that bind to 5 pMHC multimers. For each pMHC, we trained and optimized the model using TCRs from 3 donors and tested the optimized model using TCRs from the last one. As a result, DeepAIR achieved the best performance in all tested donors with a median AUC of 0.942 (Figure 4A, Table 1, and Supplementary Table 5). It is interesting to note that DeepAIR-stru achieved the second-best performance in all the tests. This reveals that structure information contributed most to the advantage of the DeepAIR in predicting the AIR-antigen binding reactivity.

Interestingly, the performance of the methods using structure information, including DeepAIR and DeepAIR-stru, appear to be much more stable than sequence-based methods, including DeepAIR-seq, TCRAI, DeepTCR, and soNNia, as evidenced by the lower value of the coefficient of variance in all tests (Figure 4B). The result also reveals that structure information indeed helps to improve the robustness of the model in predicting the AIR-antigen binding reactivity.

To investigate which part of the structure is particularly important for DeepAIR to predict the AIR-antigen binding reactivity, we highlighted the CDR3 loops of TCR with the highest DeepAIR attention weights in predicting the recognition of ELAGIGILTV (MART-1 protein, melanoma) (Figure 4C), in which DeepAIR achieved the highest AUC score (Supplementary Table 4). We note that, similar to what we observed in the prediction of binding affinity, in this task DeepAIR paid more attention to the antigen-β-chain-contacting residues and antigen-contacting residues on the α chain and β chain, respectively. This implies the distinct roles of the α and β chains in the AIR-antigen binding complex. Interestingly, although the sequences in the highlighted region are diverse (Figure 4D), they somehow constitute relatively conserved structures (Figure 4C). On the contrary, the sequences and structures are both highly diverse for the TCRs without binding reactivity (Supplementary Figure 2). This further illustrates why structured-based methods could significantly outperform sequence-based methods in this study.

To evaluate the performance of DeepAIR for predicting the BCR-antigen binding reactivity, we collected experimentally validated antigen binding BCRs from the Immune Epitope Database (IEDB)^27^. After the data curation, 553 BCRs with known binding reactivity, including 158 BCRs that bind to the envelope glycoprotein (ENV) of human immunodeficiency virus (HIV), 61 BCRs that bind to hemagglutinin (HA) of flu, 24 BCRs that bind to the circumsporozoite (CS) protein of *Plasmodium falciparum*, 21 BCRs that bind to the spike glycoprotein (GP) of Zaire ebolavirus (EBOV), were used in this study. We randomly split the BCRs into the training data (70% of the whole data), validation data (20%), and independent test data (10%) to evaluate the performance of each method for predicting the BCR binding reactivity to ENV (HIV), HA (flu), CS protein (*P. falciparum*) and GP (EBOV), respectively. As a result, DeepAIR achieved a median AUC of 0.913 in predicting the AIR-antigen binding reactivity for BCRs (Table 1 and Supplementary Table 6), outperforming all the other methods, including DeepAIR-stru (median AUC=0.722), DeepAIR-seq (median AUC=0.864), and soNNia (median AUC=0.660) (Table 1, and Supplementary Table 6). We note that structure-based DeepAIR-stru achieved a lower performance in predicting AIR-antigen binding for BCR but not in that for TCR, compared to sequence-based DeepAIR-seq. Given that the AlphaFold2-predicted BCR structure is much worse than the AlphaFold2-predicted TCR structure (Figure 2F), it implies the quality of the predicted AIR structure can greatly impact the prediction of AIR-antigen binding. On the other hand, it is interesting to note that DeepAIR still successfully extracted useful information from the predicted BCR structure to improve the prediction accuracy of AIR-antigen binding reactivity (Table 1).

### Immune repertoire classification

The immune repertoire consists of AIRs that recognize antigens associated with target disease and confounding AIRs that are irrelevant to the target antigen or the target disease^28^. To classify the immune repertoire for individuals with different health status, DeepAIR uses a two-step procedure including the receptor-level probability prediction and repertoire-level multiple instance learning (MIL, see ‘Materials and Methods’).

To evaluate the performance of DeepAIR for classifying the immune repertoire, we collected the single-cell V(D)J sequencing data for nasopharyngeal carcinoma (NPC)^28^ and inflammatory bowel disease (IBD)^28^, respectively. From the violin plots of predicted AIR (TCR and BCR) probabilities in each NPC sample and nasopharyngeal lymphatic hyperplasia (NLH) sample, we can observe that the values in the NPC sample are generally higher than those in the NLH samples, and that the constitution of AIR repertoire is diverse across samples (Figure 5A and 5B). Meanwhile, we also observe the existence of confounding AIRs from the distribution of predicted probability for each AIR from NPC samples and NLH samples. Similar observations are found in the IBD and healthy samples (Figure 5C and 5D). It is difficult to separate the disease samples from control samples using the original probabilities. However, after applying MIL, the immune repertoires for individuals with different health status could be well separated (Figure 5). This is consistent with the observation that majority-voting-based MIL has high AUC values in classifying the TCR and BCR repertoire, respectively (Table 1).

**Figure 5.**
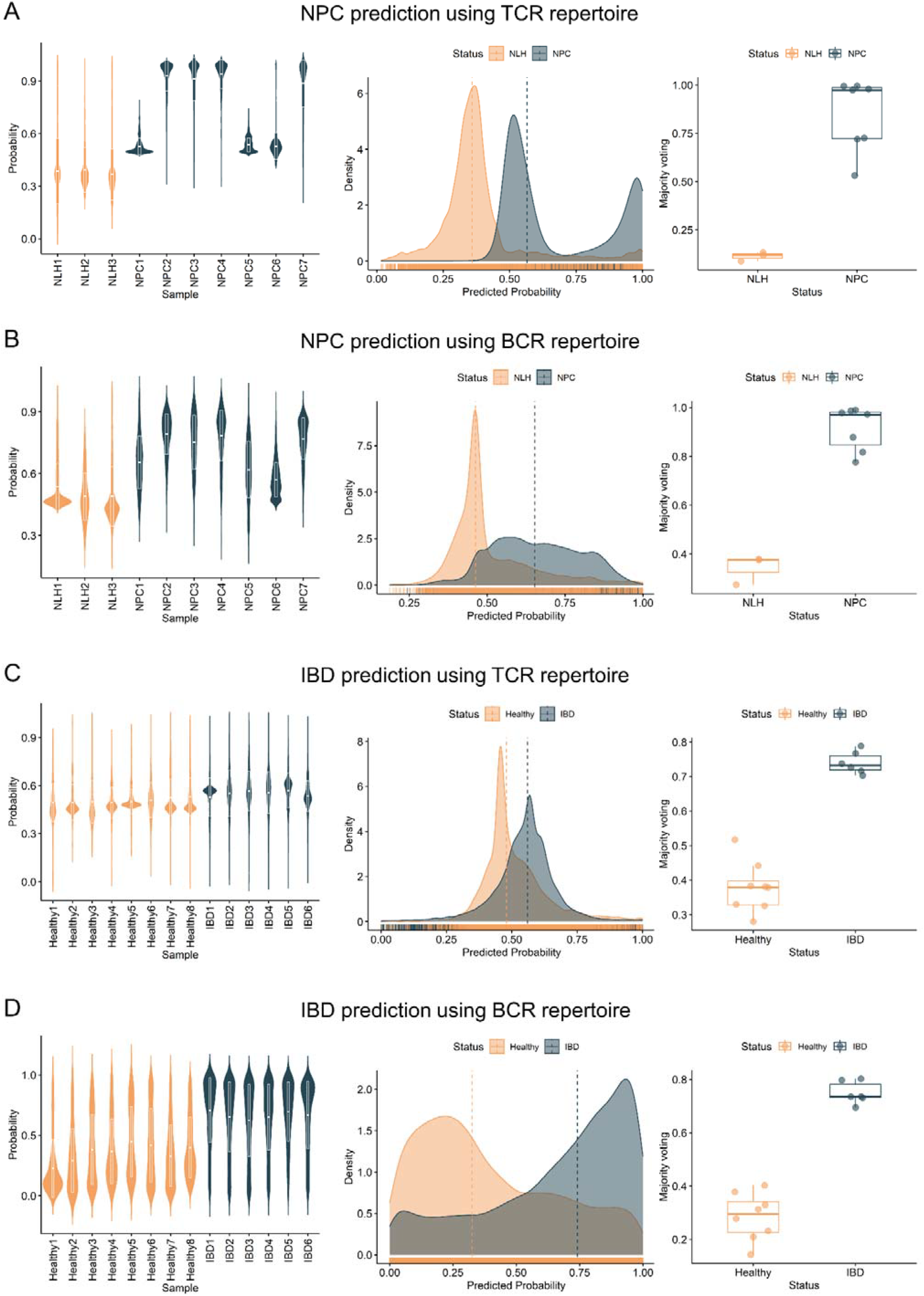
Performance of DeepAIR for the classification of the immune repertoire with nasopharyngeal carcinoma (NPC) and inflammatory bowel disease (IBD), respectively. The violin plot of the predicted AIR probability values for each sample (left), the distribution of the predicted AIR probability values for each health status (middle), and box plot of immune repertoire MIL value for each sample (right) in (**A**) the prediction of NPC using TCR repertoire, (**B**) the prediction of NPC using BCR repertoire, (**C**) the prediction of IBD using TCR repertoire, and (**D**) the prediction of IBD using BCR repertoire, respectively.

## Discussion

Building a reliable prediction model for the adaptive immune receptor (AIR)-antigen binding can significantly assist the experimental study of the adaptive immune system. Current models, such as DeepTCR, TCRAI, and soNNia, are based on the sequence information of AIR^29^. We hypothesized that integrating the structure information of AIR into the model may improve the prediction accuracy for the AIR-antigen binding. Therefore, in this study, we present DeepAIR, a deep learning framework integrating structure information for the AIR-antigen binding analysis. DeepAIR significantly outperformed sequence-based methods, including DeepTCR, TCRAI and soNNia, in predicting the AIR-antigen binding reactivity. We created two versions of DeepAIR, including structure-based DeepAIR-stru and sequence-based DeepAIR-seq, to investigate the contribution of the structure information to the performance of DeepAIR. Our experiments demonstrate that DeepAIR-stru significantly outperformed DeepTCR, TCRAI and soNNia, while DeepAIR-seq did not achieve the best prediction performance (Supplementary Tables 4 and 5). The performance comparison reveals that the integration of structure information significantly contributed to the superior performance of DeepAIR. Indeed, DeepAIR successfully captured structure patterns from antigen binding AIRs to distinguish them from others (Figure 4C).

DeepAIR uses the AIR structures predicted by AlphaFold2^30^. The major advantage of using predicted structures is that DeepAIR can analyze any AIR as long as its sequence information is available for structure prediction. This is critical for AIR analysis due to the fact that experimentally validated structures are not available for most of AIRs in the immune repertoire^31^. Moreover, AlphaFold2 demonstrated high accuracy competitive with experimental structures according to the results of 14^th^ Critical Assessment of Protein Structure Prediction (CASP14)^28^. Our analysis also showed that the median prediction accuracy for the CDR3 region of AIRs using long sequence is comparable to the median accuracy AlphaFold2 achieved in CASP14 (Supplementary Tables 2 and 3). We therefore believe that it is reliable to use AlphaFold-2 predicted structures in DeepAIR. However, the accuracy of the AlphaFold2-predicted BCR structures is much worse than that of the AlphaFold2-predicted TCR structures (Figure 2F). Consistently, DeepAIR-stru showed an inferior performance in predicting the BCR-antigen binding than in predicting the TCR-antigen binding. Therefore, the accuracy of DeepAIR-stru highly depends on the accuracy of the predicted structures. To alleviate such bias introduced by the predicted structures, DeepAIR also integrate information from sequence and gene features using the multimodal feature fusion module to jointly contribute to its prediction. The performance of DeepAIR is significantly better than both DeepAIR-stru and DeepAIR-seq (Figure 3B and 4A). It reveals that the gating-based attention and tensor fusion mechanism in the fusion module successfully extracted distinguishable features from both structures and sequences to achieve the superior performance. Moreover, although AlphaFold2-predicted BCR structures are less accurate than the predicted TCR structures, the performance of DeepAIR for BCR-antigen binding is still comparable to TCR (Table 1), suggesting the robustness and scalability of DeepAIR in AIR-antigen binding prediction.

DeepAIR is an interpretable model that shows important residues in both α and β chains that are important to the AIR-antigen binding using the attention weights. Several studies have shown the importance of β chain contact residues in AIR-antigen binding^32,33^, which can also be learned by the sequence-based deep learning model^31^. Indeed, DeepAIR can highlight important residues on the β chain that are the contact residues between β chain and antigen (Figure 3D). DeepAIR can also identify the critical residues on the α chain that contact between α chain and β chain to stabilize the AIR structure (Figure 4C). The majority of the AIR-antigen studies focused on the β chains of AIRs and their contact residues with antigens^34^. DeepAIR further enables the examination of AIR-antigen complex stabilization by highlighting both structurally and functionally important residues in both α and β chains.

There are some limitations of the current study. First, the TCR-pMHC binding affinity value used in this study is presented by the UMI count of TCRs captured by the pMHC rather than the real binding affinity^23^, as measuring the real binding affinity between TCRs and pMHCs is challenging. Although shape complementarity statistic and buried surface area (BSA) have often been used to describe the TCR-pMHC interaction, neither of them are reliable indicators of TCR-pMHC binding affinity^34^. Second, due to the limited availability of BCR-antigen binding affinity data, this study did not evaluate the performance of DeepAIR in predicting the BCR-antigen binding affinity. Since the BCRs and antibodies from the same B cell have nearly the same antigen binding affinity^35^, the prediction of BCR-antigen binding affinity may mostly equal to that of antibody-antigen binding affinity. With more data available in the future, we will combine these two tasks and investigate the prediction power of DeepAIR on antibody (BCR)-antigen binding affinity. Another limitation is that the AIR structures predicted by AlphaFold2 are unliganded (Figure 2G). However, it is known that the CDR3 loops of TCR undergo a conformational change upon pMHC binding^26^. Similar scenarios also occur in BCRs upon antigen binding^36^. The conformational changes of AIR structures can impact the AIR-antigen binding; however, such changes cannot be predicted by AlphaFold2. Novel approaches that are capable of accurately predicting the conformational changes of AIR structures upon antigen binding will undoubtedly benefit the research of AIR-antigen recognition. Meanwhile, as a generalized protein structure prediction tool, the prediction model of AlphaFold2 is not optimized for predicting the structure of AIR. In fact, AlphaFold2 showed an inferior performance in predicting the CDR3 structure for BCR than that for TCR (Figure 2F). It’s consistent with result that DeepAIR showed a worse performance in predicting binding reactivity for BCR than that for TCR. The increased accuracy of the predicted structure can greatly improve the performance of DeepAIR-like structure-based methods. Finally, for the immune repertoire that includes a high number of diverse AIRs, it is time-consuming to predict the structure of each AIR in the immune repertoire using AlphaFold2. To tackle this issue, a lighter and faster prediction model that is specifically designed and optimized for the AIR structure will greatly benefit DeepAIR and other structure-based strategies in the future.

In conclusion, DeepAIR is a comprehensive and interpretable deep-learning framework for AIR-antigen binding analysis integrating both sequence and structural information. DeepAIR shows outstanding prediction performance in terms of AIR-antigen binding reactivity and outperformed state-of-the-art predictors. We anticipate that DeepAIR may serve as a prominent tool for profiling highly antigen-interacting AIRs, thereby better informing the design of personalized immunotherapy.

## Methods

### Curation of the dataset for analysis of the AIR-antigen binding

We downloaded the pMHC-specific binding data of TCR from the 10x Genomics website (https://support.10xgenomics.com/single-cell-vdj/datasets). The dataset was then processed using the Integrative COntext-specific Normalization (ICON) workflow^29^. First, for each sample, the dataset included both single-cell RNA-seq data and paired α/β-chain single-cell TCR-seq data. We then used the single-cell RNA-seq-based quality control to remove the low-quality cells, such as doublets and dead cells. Doublets refer to the T cells with over 2500 detected genes per cell, while the cells with over 20% of mitochondrial gene expression or less than 200 detected genes per cell were considered dead cells. Then we estimated the background noise using the six negative-control dextramers, which are supposed to have no binding affinity with any of the TCRs in the dataset. The background noise threshold was assigned to each donor to remove false-positive bindings according to the signal and noise distributions. The α/β chains of the rest T cells were further checked based on single-cell TCR-seq data. For each cell, the chains with non-productive or non-high-confidence sequences were removed from the dataset. T cells with only a single chain were then removed from the dataset. If multiple α or β chains were detected in a T cell, the chain with the highest UMI counts was retained. After data curation, 38,558 paired TCR α/β chains that bind to 7 pMHC multimers, including ELAGIGILTV from the MART-1 protein of melanoma, GILGFVFTL from the M1 protein of the influenza virus (flu), KLGGALQAK from the IE1 protein of the cytomegalovirus (CMV), GLCTLVAML from the BMLF1 protein of the Epstein-Barr virus (EBV), AVFDRKSDAK from the EBNA4 protein of EBV, IVTDFSVIK from the EBNA3B protein of EBV, and RAKFKQLL from the BZLF1 protein of EBV, were used in this study (Supplementary Table 1). The observed binding affinity between TCR and pMHC was estimated by the TCR UMI counts for the specific pMHC minus the average TCR UMI counts for negative controls.

We also downloaded experimentally validated TCRs from a recent SARS-CoV-2 study^33^. The SARS-CoV-2 virus dataset contains 592 paired TCR α/β chains that bind to three pMHC multimers from the SARS-CoV-2 virus. These pMHC multimers include LTDEMIAQY and YLQPRTFLL from the spike protein, and TTDPSFLGRY from the ORF1ab polyprotein, respectively.

We downloaded experimentally validated data of BCR with a known epitope from the Immune Epitope Database (IEDB)^34^ (https://www.iedb.org/). To download the BCR data from IEDB, “B cells” were selected in “Assay”, with “Host” set to “Human” in the searching option. A total of 996 BCRs were downloaded from IEDB. The BCRs were further filtered by selecting the unique ones with paired full-length chains and known experimentally validated antigens. Meanwhile, six BCRs were removed from the dataset as AlphaFold2 failed to predict their structure. Finally, 553 BCRs were used in this study. Among them, 264 BCRs were used as the positive samples for the model, including 158 BCRs that bind to the envelope glycoprotein (ENV) of human immunodeficiency virus (HIV), 61 BCRs that bind to the hemagglutinin (HA) of flu, 24 BCRs that bind to the circumsporozoite (CS) protein in *Plasmodium falciparum*, and 21 BCRs that bind to the spike glycoprotein (GP) of Zaire ebolavirus (EBOV). The rest BCRs were then used as the negative data.

### Curation of the data for classification of the immune repertoire

We downloaded the raw single-cell V(D)J sequencing data, including RNA and TCR/BCR sequencing data for nasopharyngeal carcinoma^28^ (SRP262300) and inflammatory bowel disease^36^ (SRP181666) from the Sequence Read Archive (SRA)^37^ (https://www.ncbi.nlm.nih.gov/sra). Then we used the Cell Ranger pipeline (v6.1.2, 10x Genomics, Pleasanton, CA) to analyze the single-cell sequencing data. The FASTQ reads were aligned to the GRCh38 human reference (v5.0.0) to extract the gene expression matrix and TCR/BCR sequences for each cell. In each cell, the chains with non-productive or non-high-confidence sequences were removed from the dataset. If multiple α or β chains were detected in a T cell, or multiple heavy or light chains were detected in a B cell, the chain with the highest UMI counts was retained for that cell. Those T cells and B cells that had only a single chain, with over 5000 or less than 200 detected genes per cell, or with over 20% of mitochondrial gene expression, were further removed from the dataset. After the data curation process, 18,985 paired TCR α/β chains and 15,539 paired BCR heavy/light chains from 6 patients with NPC and 3 patients with nasopharyngeal lymphatic hyperplasia (NLH) were obtained. While for IBD, 38,316 paired TCR α/β chains and 27,877 paired BCR heavy/light chains from 6 patients and 8 healthy controls were processed for further analysis.

### Prediction of the AIR structure

We used amino acid sequences of the paired chains (i.e., the α and β chains for TCRs, or heavy and light chains for BCRs) as the input to AlphaFold2^13^ to predict AIR structures. Then the structure of the CDR3 loop was extracted from each predicted AIR structure. Specifically, multiple sequence alignments (MSAs) for the CDR sequences were generated by HHBlits^38^ with the following command: hhblits -i <input-file> -o <result-file> -oa3m <result-alignment> -n 3 -e 0.001 -d <uniclust30>. HHBlits searches the sequences with three iterations against the consensus sequences in the uniclust30 database, clustering the UniProtKB^39^ sequences at the level of 30% pairwise sequence identity^40^. We accepted MSA hits with an e-value of lower than 0.001. HHsearch^41^ was used to identify the top 20 ranked templates through a clustered version of the PDB70^21^, which contains PSI-BLAST^42^ alignments produced with sequences of PDB full chain representatives (<70% sequence identity) as queries. The accepted templates and MSAs were used as the input features for AlphaFold2^42^ (version v2.1.1). Specifically, the monomer predicted TM-score (pTM) model, which is the original CASP14 model fine-tuned with the pTM layer, provides a pairwise confidence measure and therefore was used for the structure prediction.

### The construction of the DeepAIR framework

#### Overview

DeepAIR was designed with a feature encoding backbone and multiple task-specific prediction layers for addressing both receptor-level analysis tasks, including binding affinity prediction (DeepAIR with a main regression layer) and binding reactivity prediction (DeepAIR with a classification layer), and repertoire-level analysis tasks (DeepAIR with a multiple instance learning (MIL) layer) such as the repertoire classification (e.g., the disease diagnosis based on the adaptive immune repertoire) (Figure 1D). The feature encoding backbone of DeepAIR consists of a multi-channel feature extraction module and a multimodal feature fusion module. In the multi-channel feature extraction module, three feature encoders are involved, i.e., V(D)J gene encoder, sequence encoder, and structure encoder. The V(D)J gene encoder embeds the V(D)J gene segment information via a trainable embedding layer. The sequence encoder generates a high-level representation of sequence information for the two TCR/BCR chains (CDR3 regions) based on a multi-layer transformer encoder. The structure encoder utilizes the pre-trained AlphaFold2 to extract initial structure-information embedded features and then employs a two-layer one-dimensional convolution module to fine-tune and recalibrate these features specifically for better AIR-antigen binding prediction. The multi-channel feature extraction module is followed by the multimodal feature fusion module to integrate the obtained features obtained from feature extracting channels via a gating-based attention mechanism^43^ as well as a tensor fusion^44^ for generating a comprehensive representation of the AIR receptor. Then, the task-specific prediction layers map the obtained receptor representation to the predicted results (Figure 1D).

#### Immune-receptor-level analysis

##### Binding affinity prediction

During the binding affinity prediction, the resulting features obtained after the feature encoding backbone were input into a regression layer (composed of a multilayer perceptron [MLP]), which serves to map its input to the binding affinity prediction of adaptive immune receptors (TCR/BCR). In the training phase, a multi-task training strategy with adding an auxiliary affinity grading layer was utilized to train DeepAIR for the binding affinity prediction. Two training loss functions were utilized. The primary loss function was the mean squared error (MSE) loss for the main regression layer, which encourages DeepAIR to directly predict an accuracy affinity score. The auxiliary loss is the categorical cross-entropy (CE) loss for the auxiliary affinity grading layer, which encourages DeepAIR to learn the accuracy affinity orders (i.e., receptors with higher binding affinity have higher affinity grades). Specifically, the MSE loss *ℒ*_*MSE*_ and CE loss *ℒ*_*CE*_ are defined as:

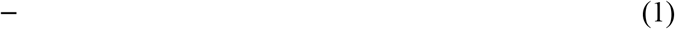

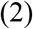

where *N* is the sample number, *y* is the ground truth binding affinity, *ŷ*_*i*_ denotes the predicted binding affinity,*C* represents the number of stages, 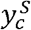 is the ground truth affinity stage, and *p*_*c*_ is the probability for the *c*^*th*^ stage. The total loss of DeepAIR binding affinity prediction, *L*_*BAP*_ is defined as:

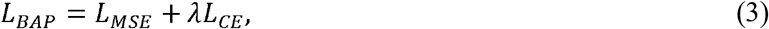

where *λ* is the hyper-parameter to adjust the influence of the auxiliary loss, which was set to 1 in this work.

##### Binding reactivity prediction

Similarly, for binding reactivity prediction, DeepAIR utilizes an MLP classification layer to map the embedded features after the feature encoding backbone to the prediction of the AIR-antigen binding reactivity, i.e., identifying which antigen epitope an assessed AIR (BCR/TCR) can bind to. During the training of the DeepAIR for the AIR-antigen binding reactivity prediction, the above-mentioned categorical cross-entropy (CE) loss is used to allow DeepAIR to learn informative features.

#### Immune-repertoire-level analysis

In the repertoire-level analysis tasks, the characteristics of the entire immune repertoire with massive receptors or the associations between the entire immune repertoire and an interesting subject-level status such as disease or healthy are evaluated. Different from receptor-level analysis tasks, where each receptor has the information to conduct the prediction, the repertoire-level analysis task needs to comprehensively integrate the information of all possible related receptors in an immune repertoire to make a prediction. The repertoire-level analysis task can be formulated as a typical multiple instance learning (MIL) problem^45^, where each repertoire can be regarded as a bag containing receptors that are instances. Note that it is challenging as there exist massive instances in a repertoire and only a fraction of these receptors are correlated with the interested subject-level status and therefore are discriminative.

In this work, we focused on the repertoire classification, which is a kind of repertoire-level analysis task when the prediction output variable is a category. DeepAIR uses a two-stage pipeline to address this MIL task. At the first stage, we trained the DeepAIR with a classification layer to obtain receptor-level predictions, i.e., the category probabilities referring to each AIR. Then DeepAIR comprehensively summarizes all receptor-level predictions with a majority-voting based MIL layer to perform the entire-repertoire-level prediction.

Specifically, assuming *R* is a repertoire with *M* receptors *{r*_1_, *r*_2_, …, *r*_*M*_*}*, DeepAIR first trains a receptor-level prediction model *f(r*_*m*_, *θ*) with the classification layer to predict the repertoire-level-category probability as *p*_*m*_ = *f*(*r*_*m*_, *θ*), where *θ* represents the model’s parameters. Then DeepAIR uses the MIL layer to integrate the predictions (votes) of all receptors to predict the repertoire-level-category probability of the label *Y* with a transformation *ϕ*_*MV*_ given by

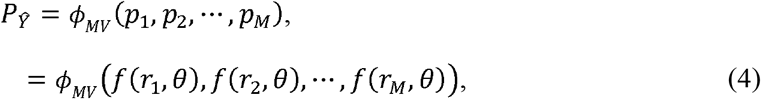

where *ϕ*_*MV*_ represents a majority-voting strategy and is defined as:

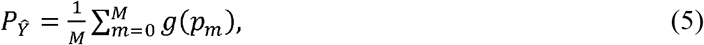

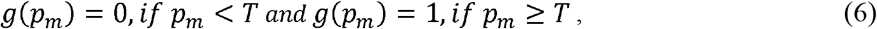

where *T* is a threshold that has been set to 0.5 in this study. In this work, we evaluated the performance of DeepAIR (with MIL layers) on two repertoire classification tasks including the diagnoses of the nasopharyngeal carcinoma (NPC)^28^ and inflammatory bowel disease (IBD)^41^.

### Comparison of different methods for the immune-receptor-level analysis

#### Predicting TCR-antigen binding affinity and reactivity with DeepTCR

The TCR-antigen binding reactivity and affinity prediction using DeepTCR was performed by following the instructions provided in the study by Sidhom *et al*^42^. For each TCR, we used the single paired α and β TCR chains, with CDR3 amino acid sequence and V(D)J gene usage as the input to DeepTCR. DeepTCR encodes the amino acids to the numbers between 0 and 19 and uses categorical variables to represent the genes in the V(D)J gene usage in the feature calculation step. DeepTCR then implements a variational autoencoder to transform the features into a latent space that is parametrized by a multidimensional unit Gaussian distribution^7^. To cluster the antigen binding TCR sequences, a Euclidean distance in the latent space was used to measure the closeness between any two TCR sequences. To predict the binding affinity, a supervised TCR sequence regression was performed with the UMI (unique molecular identifier) counts as the measure for the predicted binding affinity.

#### Predicting TCR-antigen binding reactivity with TCRAI

We used the paired α and β TCR chains, with CDR3 amino acid sequence and V, J genes for each chain, as the inputs^44^. For each CDR3 sequence, TCRAI applies the one-hot representation scheme to generate an integer vector for the given CDR3 sequence. For the V and J genes, TCRAI encodes the V and J gene seperately^45^. Then TCRAI builds a convolutional neural network architecture to process the input information and provides a prediction for the binding reactivity.

#### Predicting TCR-antigen binding reactivity with soNNia

We used the paired receptor chains (i.e., the α and β chains for TCRs, the heavy and light chains for BCRs, respectively), with the CDR3 amino acid sequence and V, J genes for each chain, as the inputs^46^. soNNia divides the sequence features into three categories: V(D)J gene usage, CDR3 length, and CDR3 amino acid composition. The inputs from each category are first propagated through the neural network model, and then are combined and transformed through a dense layer of the deep neural network. A log-likelihood ratio is then computed as a functional classifier for the binding reactivity prediction.

### Statistical analysis

The AUC values of the ROC curves were calculated by the pROC R package^47^. The statistical significance of the AUC differences was determined by the paired Wilcoxon test^48^. The statistical significance of ROC differences was determined by the Delong method^49^. All *p*-values are two-sided unless stated otherwise. The *p*-value of less than 0.05 was defined as being statistically significant.

## Data availability

All data needed to evaluate the conclusions in the paper are presented in the paper and/or the Supplementary Materials. The AIR data, source code and well-trained DeepAIR models are available at https://github.com/TencentAILabHealthcare/DeepAIR

## Supplementary materials

**Supplementary Figure 1.**
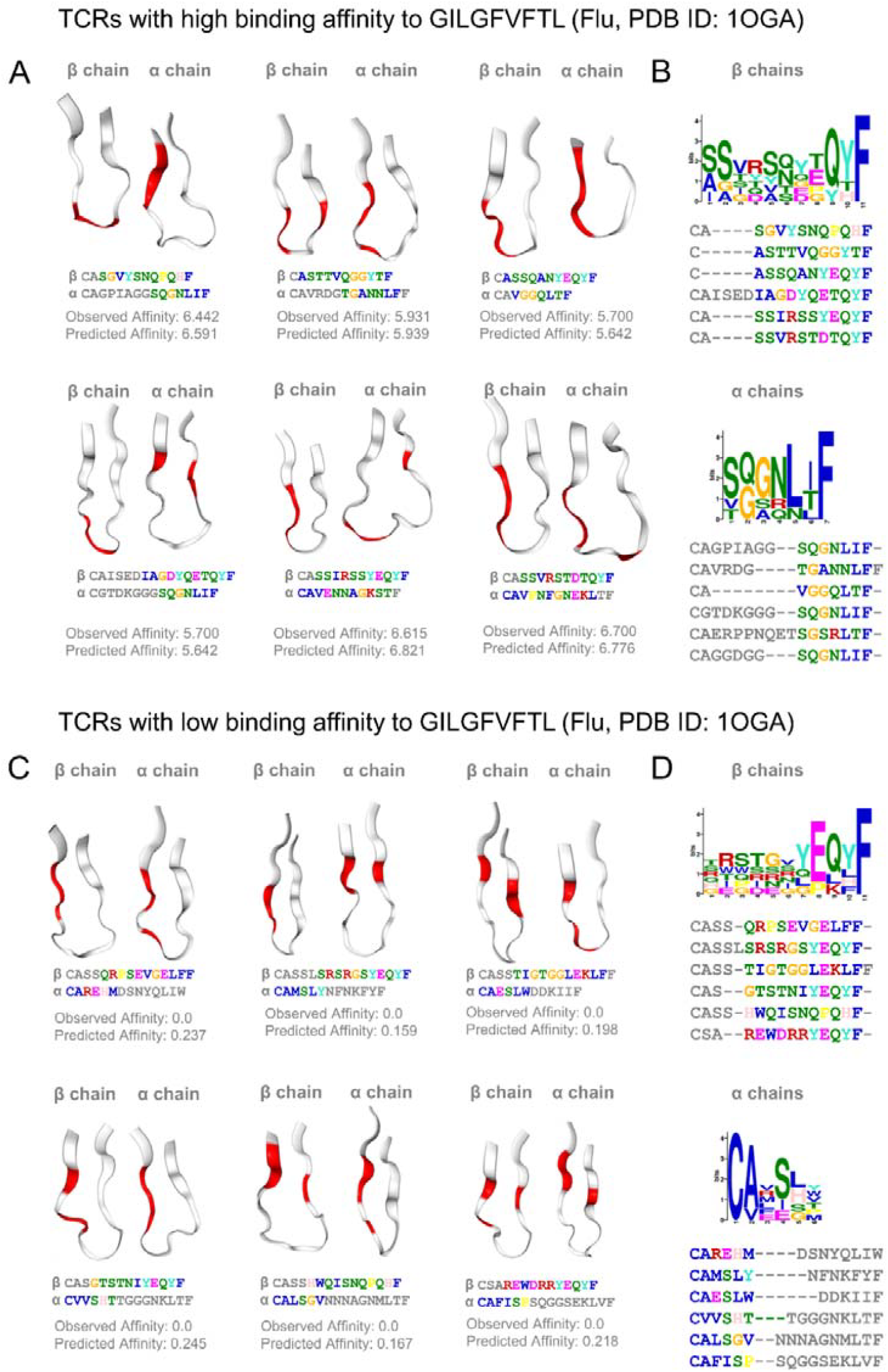
The structures and sequences of TCRs that have high and low binding affinity to GILGFVFTL (M1 protein, flu). (**A**) DeepAIR highlights similar partial structures in TCRs with high binding affinity to GILGFVFTL. (**B**) The sequences of TCRs in the subfigure (A) are quite diverse. There are no similar (**C**) partial structures or (**D**) sequences observed in TCRs with low binding affinity to GILGFVFTL. The most salient regions informative for DeepAIR to make the correct prediction are highlighted in red.

**Supplementary Figure 2.**
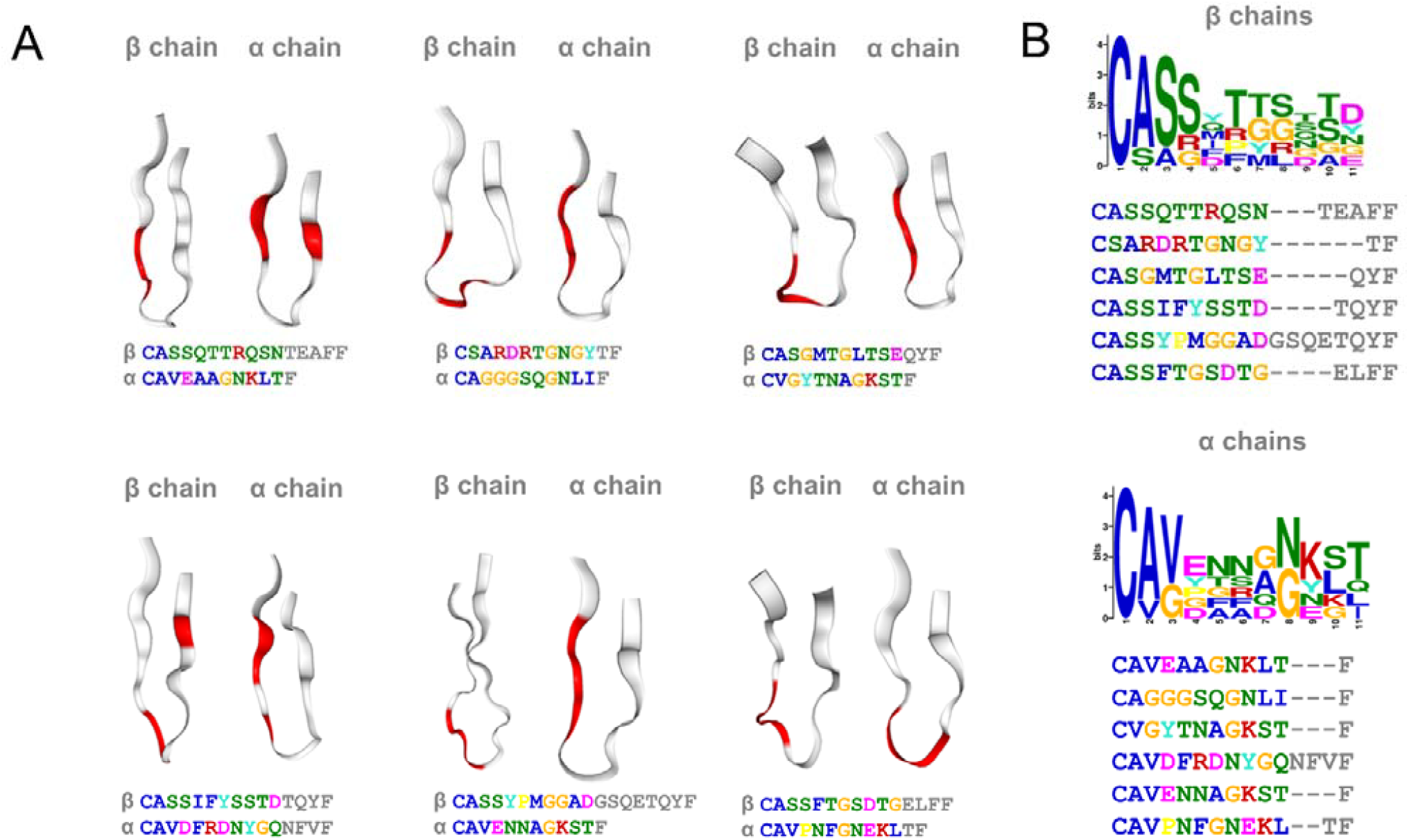
The structures and sequences of TCRs that have no binding reactivity to ELAGIGILTV (MART-1 protein, melanoma). (**A**) The structures of TCRs without the binding to ELAGIGILTV are illustrated, and the most salient regions informative for DeepAIR to make the correct prediction are highlighted in red. (**B**) The sequences and recognized motifs of the corresponding beta and alpha chains of the TCRs in the subfigure (C) Both the sequences and structures of these TCRs are quite diverse.

**Supplementary Table 1.**
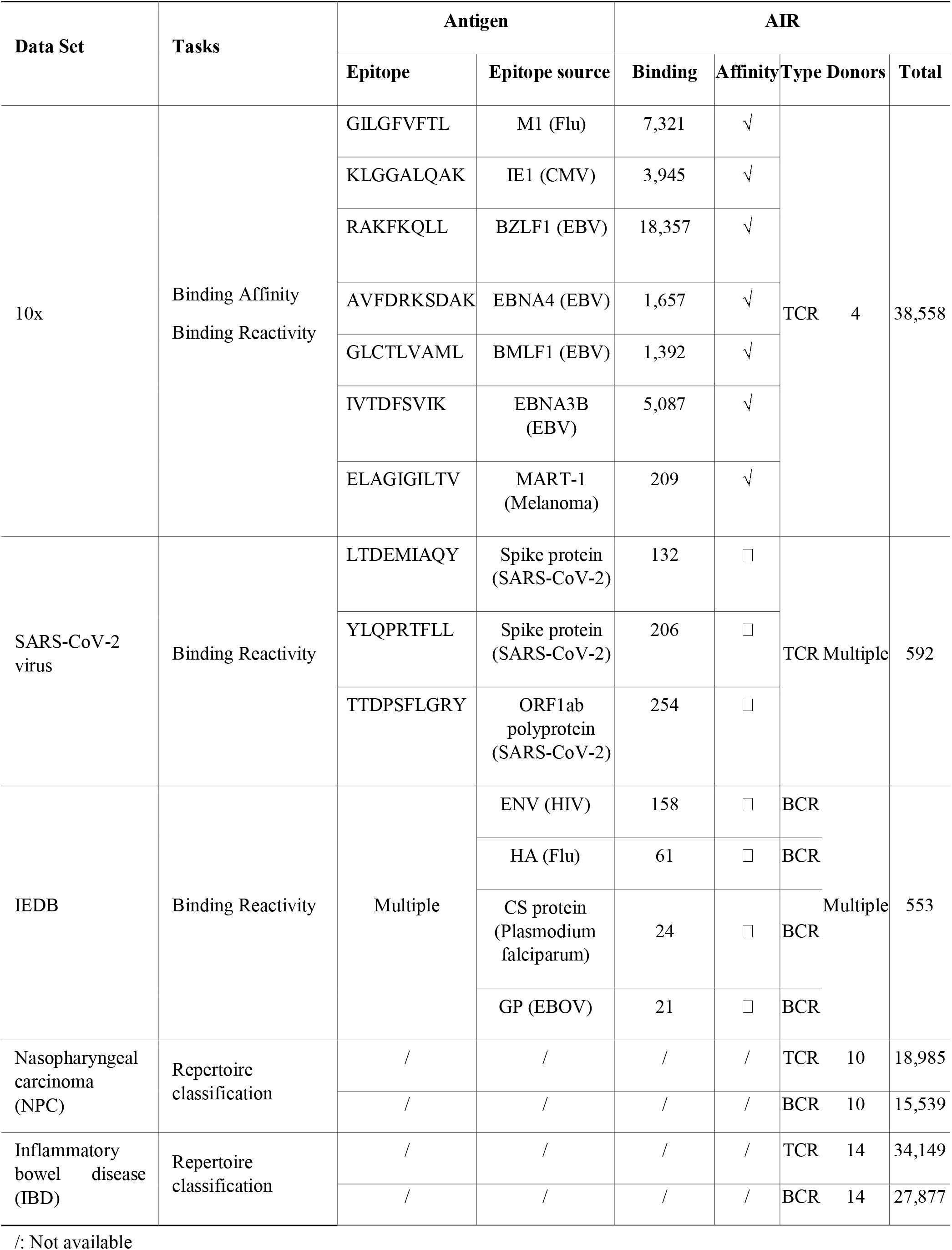
Statistical summary of the datasets curated in this study.

**Supplementary Table 2.** RMSDs between the predicted and experimentally validated TCR structures.

| TCR status | PDB | Long Sequence |  |  | Short sequence |
| --- | --- | --- | --- | --- | --- |
|  |  | TCR Beta Chain | CDR | CDR3 | CDR3 |
| Free TCR | 1KGC | 0.601 | 0.371 | 0.233 | 6.053 |
|  | 2VLM | 1.533 | 0.45 | 0.83 | 7.741 |
|  | 7N1C | 1.11 | 0.199 | 0.333 | 12.426 |
|  | 7N1D | 0.519 | 0.442 | 0.184 | 7.446 |
|  | 7EA6 | 0.875 | 0.304 | 0.309 | 8.767 |
|  | 6VTH | 0.759 | 0.384 | 0.362 | 6.275 |
| TCR in Binding Complex | 1OGA | 1.306 | 0.337 | 0.294 | 8.429 |
|  | 6V0Y | 0.948 | 0.337 | 2.192 | 11.87 |
|  | 1AO7 | 0.629 | 0.411 | 2.034 | 12.458 |
|  | 3O4L | 0.565 | 0.615 | 1.339 | 2.107 |
|  | 5HHM | 0.601 | 0.304 | 0.205 | 9.8 |
|  | 3QH3 | 0.757 | 0.307 | 0.966 | 12.406 |

**Supplementary Table 3.** RMSDs between the predicted and experimentally validated BCR structures.

| BCR status | PDB | Long Sequence |  |  | Short sequence |
| --- | --- | --- | --- | --- | --- |
|  |  | BCR Heavy Chain | CDR | CDR3 | CDR3 |
| Free BCR | 3UJI | 4.974 | 0.314 | 0.962 | 13.183 |
|  | 5CSZ | 1.338 | 0.297 | 0.601 | 14.675 |
|  | 6D01 | 2.712 | 0.371 | 2.345 | 13.258 |
|  | 6AZM | 0.731 | 0.554 | 3.081 | 11.565 |
|  | 5BK0 | 3.317 | 1.368 | 3.219 | 10.541 |
|  | 4G6F | 3.175 | 1.716 | 1.504 | 1.521 |
| BCR in Binding Complex | 6ZTD | 1.173 | 0.653 | 0.533 | 4.287 |
|  | 5IFH | 1.191 | 0.479 | 1.316 | 3.85 |
|  | 4EVN | 1.512 | 0.718 | 0.377 | 8.942 |
|  | 6Q1J | 1.631 | 3.337 | 3.225 | 7.455 |
|  | 6VU2 | 6.817 | 1.14 | 3.187 | 6.058 |
|  | 5DRW | 1.777 | 1.476 | 3.636 | 6.339 |

**Supplementary Table 4.** Performance of the binding-reactivity prediction methods on the independent test data.

| Antigen |  | AUC |  |  |  |  |  |
| --- | --- | --- | --- | --- | --- | --- | --- |
| Epitope | Epitope source | DeepAIR | DeepAIR-stru | DeepAIR-seq | TCRAI | DeepTCR | soNNia |
| LTDEMIAQY | Spike protein (SARS-CoV-2) | **0.752** | 0.681 | 0.697 | 0.623 | 0.694 | 0.612 |
| TTDPSFLGRY | ORF1ab polyprotein (SARS-CoV-2) | **0.824** | 0.776 | 0.785 | 0.755 | 0.814 | 0.781 |
| YLQPRTFLL | Spike protein (SARS-CoV-2) | **0.885** | 0.767 | 0.846 | 0.756 | 0.801 | 0.783 |
| AVFDRKSDAK | EBNA4 (EBV) | **0.881** | 0.738 | 0.599 | 0.647 | 0.675 | 0.845 |
| GILGFVFTL | BMLF1 (EBV) | **0.956** | 0.941 | 0.935 | 0.940 | 0.931 | 0.925 |
| IVTDFSVIK | EBNA3B (EBV) | **0.922** | 0.885 | 0.807 | 0.831 | 0.848 | 0.693 |
| RAKFKQLL | BZLF1 (EBV) | **0.934** | 0.907 | 0.879 | 0.933 | 0.912 | 0.860 |
| GLCTLVAML | M1 (Flu) | **0.975** | 0.890 | 0.917 | 0.915 | 0.843 | 0.751 |
| ELAGIGILTV | MART-1 (Melanoma) | 0.983 | 0.938 | 0.960 | **0.988** | 0.986 | 0.844 |
| KLGGALQAK | IE1 (CMV) | **0.872** | 0.860 | 0.770 | 0.855 | 0.853 | 0.674 |
| Median |  | **0.904** | 0.873 | 0.827 | 0.843 | 0.846 | 0.782 |

**Supplementary Table 5.**
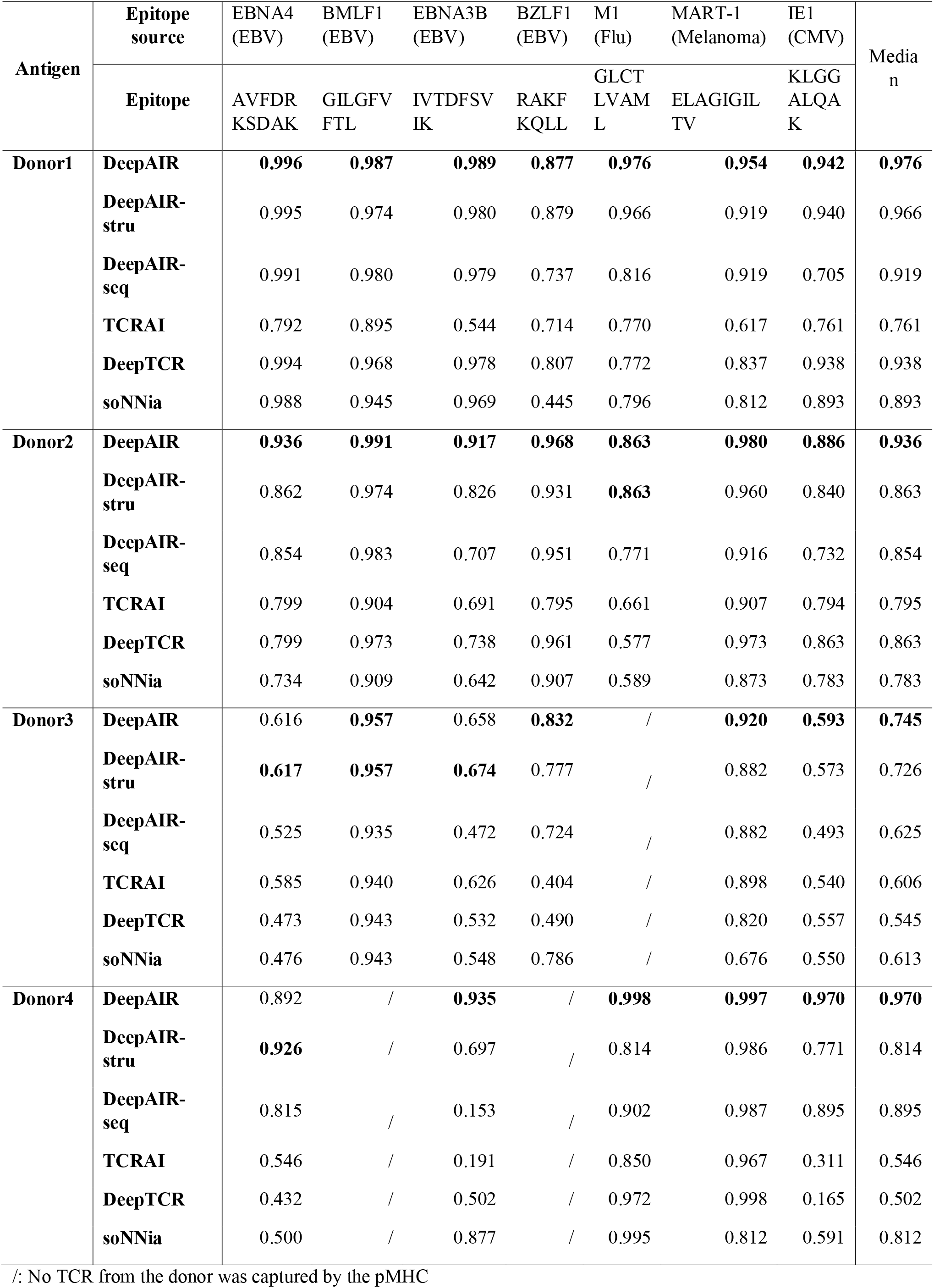
Performance of the binding-reactivity prediction methods on the leave-one-out test.

| Antigen | Epitope source | EBNA4 (EBV) | BMLF1 (EBV) | EBNA3B (EBV) | BZLF1 (EBV) | M1 (Flu) | MART-1 (Melanoma) | IE1 (CMV) | Median |
| --- | --- | --- | --- | --- | --- | --- | --- | --- | --- |
|  | Epitope | AVFDR KSDAK | GILGFV FTL | IVTDFSV IK | RAKF KQLL | GLCT LVAM L | ELAGIGIL TV | KLGG ALQAK |  |
| Donor1 | DeepAIR | **0.996** | **0.987** | **0.989** | **0.877** | **0.976** | **0.954** | **0.942** | **0.976** |
|  | DeepAIR-stru | 0.995 | 0.974 | 0.980 | 0.879 | 0.966 | 0.919 | 0.940 | 0.966 |
|  | DeepAIR-seq | 0.991 | 0.980 | 0.979 | 0.737 | 0.816 | 0.919 | 0.705 | 0.919 |
|  | TCRAI | 0.792 | 0.895 | 0.544 | 0.714 | 0.770 | 0.617 | 0.761 | 0.761 |
|  | DeepTCR | 0.994 | 0.968 | 0.978 | 0.807 | 0.772 | 0.837 | 0.938 | 0.938 |
|  | soNNia | 0.988 | 0.945 | 0.969 | 0.445 | 0.796 | 0.812 | 0.893 | 0.893 |
| Donor2 | DeepAIR | **0.936** | **0.991** | **0.917** | **0.968** | **0.863** | **0.980** | **0.886** | **0.936** |
|  | DeepAIR-stru | 0.862 | 0.974 | 0.826 | 0.931 | **0.863** | 0.960 | 0.840 | 0.863 |
|  | DeepAIR-seq | 0.854 | 0.983 | 0.707 | 0.951 | 0.771 | 0.916 | 0.732 | 0.854 |
|  | TCRAI | 0.799 | 0.904 | 0.691 | 0.795 | 0.661 | 0.907 | 0.794 | 0.795 |
|  | DeepTCR | 0.799 | 0.973 | 0.738 | 0.961 | 0.577 | 0.973 | 0.863 | 0.863 |
|  | soNNia | 0.734 | 0.909 | 0.642 | 0.907 | 0.589 | 0.873 | 0.783 | 0.783 |
| Donor3 | DeepAIR | 0.616 | **0.957** | 0.658 | **0.832** | / | **0.920** | **0.593** | **0.745** |
|  | DeepAIR-stru | **0.617** | **0.957** | **0.674** | 0.777 | / | 0.882 | 0.573 | 0.726 |
|  | DeepAIR-seq | 0.525 | 0.935 | 0.472 | 0.724 | / | 0.882 | 0.493 | 0.625 |
|  | TCRAI | 0.585 | 0.940 | 0.626 | 0.404 | / | 0.898 | 0.540 | 0.606 |
|  | DeepTCR | 0.473 | 0.943 | 0.532 | 0.490 | / | 0.820 | 0.557 | 0.545 |
|  | soNNia | 0.476 | 0.943 | 0.548 | 0.786 | / | 0.676 | 0.550 | 0.613 |
| Donor4 | DeepAIR | 0.892 | / | **0.935** | / | **0.998** | **0.997** | **0.970** | **0.970** |
|  | DeepAIR-stru | **0.926** | / | 0.697 | / | 0.814 | 0.986 | 0.771 | 0.814 |
|  | DeepAIR-seq | 0.815 | / | 0.153 | / | 0.902 | 0.987 | 0.895 | 0.895 |
|  | TCRAI | 0.546 | / | 0.191 | / | 0.850 | 0.967 | 0.311 | 0.546 |
|  | DeepTCR | 0.432 | / | 0.502 | / | 0.972 | 0.998 | 0.165 | 0.502 |
|  | soNNia | 0.500 | / | 0.877 | / | 0.995 | 0.812 | 0.591 | 0.812 |

/: No TCR from the donor was captured by the pMHC

**Supplementary Table 6.** Performance of the BCR binding-reactivity prediction methods.

| Antigen | AUC |  |  |  |
| --- | --- | --- | --- | --- |
|  | DeepAIR | DeepAIR-stru | DeepAIR-seq | soNNia |
| ENV (HIV) | **0.928** | 0.638 | 0.914 | 0.805 |
| HA (Flu) | **0.833** | 0.740 | 0.773 | 0.523 |
| CS protein (*Plasmodium falciparum*) | **0.898** | 0.722 | 0.814 | 0.519 |
| GP (EBOV) | **0.991** | 0.722 | 0.981 | 0.796 |
| Median | **0.913** | 0.722 | 0.864 | 0.660 |

